# Modularity of the metabolic gene network as a prognostic biomarker for hepatocellular carcinoma

**DOI:** 10.1101/224998

**Authors:** Fengdan Ye, Dongya Jia, Mingyang Lu, Herbert Levine, Michael W Deem

**Affiliations:** Department of Physics & Astronomy, Rice University, Houston, TX 77005, USA; Center for Theoretical Biological Physics, Rice University, Houston, TX 77005, USA; Program in Systems, Synthetic and Physical Biology, Rice University, Houston, TX 77005, USA; The Jackson Laboratory, Bar Harbor, ME 04609, USA; Department of Bioengineering, Rice University, Houston, TX 77005, USA; Department of Biosciences, Rice University, Houston, TX 77005, USA

**Keywords:** modularity, metabolism, hepatocellular carcinoma, HCC, prognosis

## Abstract

Abnormal metabolism is an emerging hallmark of cancer. Cancer cells utilize both aerobic glycolysis and oxidative phosphorylation (OXPHOS) for energy production and biomass synthesis. Understanding the metabolic reprogramming in cancer can help design therapies to target metabolism and thereby to improve prognosis. We have previously argued that more malignant tumors are usually characterized by a more modular expression pattern of cancer-associated genes. In this work, we analyzed the expression patterns of metabolism genes in terms of modularity for 371 hepatocellular carcinoma (HCC) samples from the Cancer Genome Atlas (TCGA). We found that higher modularity significantly correlated with glycolytic phenotype, later tumor stages, higher metastatic potential, and cancer recurrence, all of which contributed to poorer prognosis. Among patients with recurred tumor, we found the correlation of higher modularity with worse prognosis during early to mid-progression. Furthermore, we developed metrics to calculate individual modularity, which was shown to be predictive of cancer recurrence and patients’ survival and therefore may serve as a prognostic biomarker. Our overall conclusion is that more aggressive HCC tumors, as judged by decreased host survival probability, had more modular expression patterns of metabolic genes. These results may be used to identify cancer driver genes and for drug design.

## Introduction

Hepatocellular carcinoma (HCC) is a primary malignancy of the liver, with average survival time between 6 to 20 months without any intervention [1]. It is also the third leading cause of cancer mortality worldwide [2]. The prognosis for HCC patients remains poor [3]. Diagnosis of HCC is usually based on biomarkers, such as AFP (alpha-fetoprotein) and miR-21 [4]. However, HCC can result from a variety of risk factors, such as hepatitis B/C virus or alcoholic liver disease [5], which makes it difficult to characterize HCC with single gene biomarkers. One key to a further breakthrough in HCC therapy lies in better understanding the underlying mechanism of HCC progression.

In recent years, a significant amount of research has gone into analyzing cancer-associated pathways and networks to gain insight into the complex biological systems underlying tumor progression [6, 7]. One promising approach for breast cancer and leukemia patients has been to identify the varying patterns of cancer-associated gene expression to predict prognosis [8, 9]. In both examples, the level of organization of the cancer-associated gene network, as measured by the cophenetic correlation coefficient (CCC), was shown to be correlated with cancer risk, progression and prognosis. Inspired by these works, we here aim to characterize HCC progression and patient survival by analyzing the structure of the cancer-associated gene network in HCC.

Liver is an organ in which metabolism plays a key role. And abnormal metabolism is a hallmark of cancer [10, 11]. Therefore, we chose to analyze the expression patterns of metabolic genes in HCC patients. Unlike normal cells, cancer cells use glycolysis for energy production irrespective of the availability of oxygen, a process that is referred to as the Warburg effect or aerobic glycolysis [12, 13]. Interestingly, although aerobic glycolysis has been regarded as the dominant metabolism phenotype in cancer, recent experimental evidence shows that mitochondria are actively functional in cancer cells [14-16], and oxidative phosphorylation (OXPHOS) can enhance metastasis in certain scenarios [17, 18]. Study of the interplay between glycolysis and OXPHOS will deepen our understanding of cancer metabolism and metastasis.

To quantify the activities of the two main metabolism phenotypes in HCC, OXPHOS and glycolysis, Yu *et al.* [19] developed the AMPK and HIF-1 signatures by evaluating the expression of the downstream genes of AMPK (5’ AMP-activated protein kinase) and HIF-1 (hypoxia-inducible factor 1), in total 33 AMPK downstream genes and 23 HIF-1 downstream genes. The AMPK and HIF-1 signatures have been shown to capture the highly significant metabolic features of HCC samples [19]. In addition, the AMPK and HIF-1 signatures can associate the metabolism phenotypes of HCC samples with oncogene activities, such as MYC, c-SRC and RAS, which further validates the use of the AMPK and HIF-1 signatures in characterizing the metabolic activity of HCC samples [19]. Based on these arguments, the AMPK and HIF-1 downstream genes were chosen for the present study as a relevant set of cancer-associated genes for HCC. The strong anti-correlation between AMPK and HIF-1 activities in HCC [19] suggests the expression of these metabolic genes is modular, with the AMPK and HIF-1 downstream gene subsets as two likely modules (Fig. 1A).

**Figure 1.**
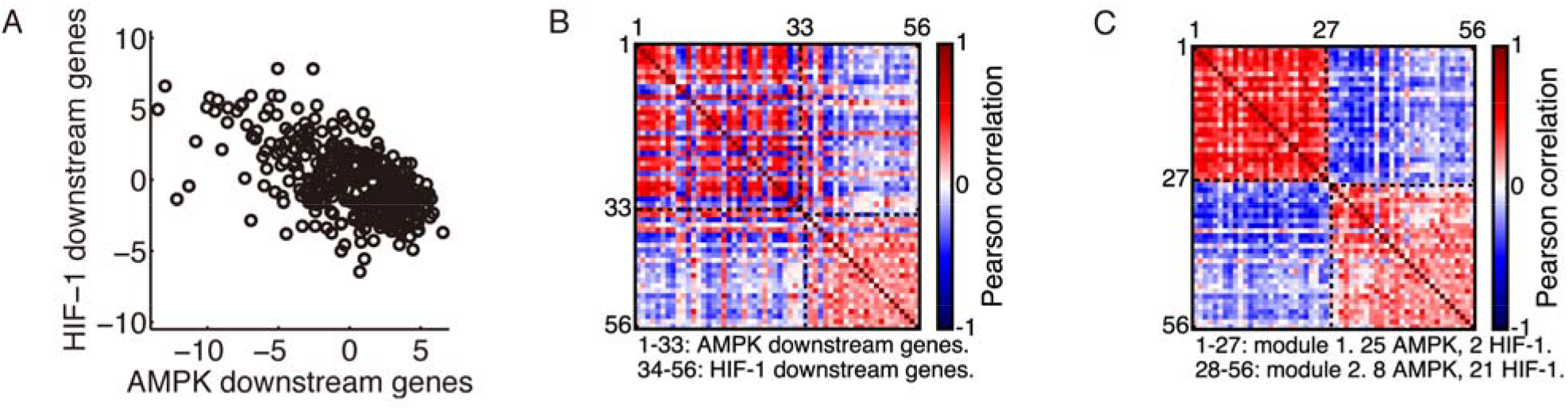
Modular gene expression pattern of the metabolic genes. **(A)** Evaluation of the AMPK and HIF-1 activities in HCC patients’ samples (*n* = 371,*r* = –0.59,*p* < 0.0001). Each point represents the AMPK and HIF-1 activities of one sample. **(B)** Correlation matrix of the 33 AMPK downstream genes and 23 HIF-1 downstream genes. **(C)** Rearranged correlation matrix calculated from the complete dataset of 371 HCC patients by the Newman algorithm. The Newman algorithm obtained a partition into two modules. Modules are labeled by black dashed lines. The red dashed lines in **(B)** and **(C)** are the diagonal elements of the correlation matrix. The red color corresponds to a correlation coefficient of 1, as each gene is fully correlated with itself. In modularity calculation, the diagonal elements were set to 0, as it was assumed that there were no self-loops.

Community structure of a gene network conveys information regarding the interaction between genes. In particular, genes within the same community cooperate much more with each other than with those in other communities. Here we utilize modularity to quantify the community structure of the metabolic gene network in HCC. Modularity is a measure of intracommunity connection strength compared to what is expected from randomly distributed connections [20, 21]. In the current context, it quantifies the ability of tumor cells to organize individual cancer-associated genes so as to maximize network efficiency. Modularity is present in almost all biological systems, from molecular interactions to macroscopic food webs [22, 23]. A general theory regarding modularity shows that high modularity systems afford greater evolutionary fitness in high stress environments or over shorter time scales, whereas low modularity systems afford greater fitness in low stress environments or over longer time scales [24, 25]. This general principle can be applied to understand the relation between modularity of cancer-related gene networks and the aggressiveness of cancer [8, 9]. Using this theory, we predict that tumors with a more modular expression pattern of cancer-associated genes, organized to counteract host defenses, are more fit and aggressive. At longer time scales, tumor growth overcomes host defenses and loses its sensitivity to host actions, and modularity is predicted to decline.

In this work, we analyzed the change of the modular expression pattern of the AMPK and HIF-1 downstream genes in HCC samples as a function of metabolism phenotypes, tumor stages, metastatic potentials and tumor recurrence. We found that (i) HCC samples with a glycolysis phenotype show significantly higher modularity than samples with an OXPHOS phenotype; (ii) HCC samples at tumor stages II-IV have significantly higher modularity than samples at stage I; (iii) HCC samples with higher metastatic potential maintain significantly higher modularity than samples with lower metastatic potential; and (iv) patients that have recurrence within 12, 24 or 36 months have significantly higher modularity than those with no recurrence within the same amount of time. These results confirm the theoretical prediction that more aggressive tumors correspond to a more modular interaction pattern of the cancer-associated gene network. We also found that modularity increases with tumor progression up to 8 months before recurrence, but then decreases. This result is examined in detail in the ‘Discussion’ section, and indicates that modularity is no longer selected for at very late stages of tumor progression. This result is also in accord with the aforementioned theoretical expectations. We further developed metrics to calculate individual modularity, which is proved to be predictive of recurrence and survival for individual HCC patients. Possible applications of modularity in terms of drug design and identifying cancer-related genes will be discussed in the ‘Discussion’ section.

## Results

To construct our HCC cancer-associated gene network, we took the 33 AMPK downstream genes and 23 HIF-1 downstream genes identified by Yu *et al.* [19] as nodes in the network. For each group of patients, the interaction patterns between genes were calculated using Pearson correlation. Simply put, two genes have a strong interaction if they show a similar trend of gene expression changing across patients. That is, if one gene expression increases and another gene expression also increases, then these genes are cooperating and strongly interacting with each other. After nodes and links were established, we applied the Newman algorithm [21] to obtain the community structure of the gene network and the corresponding modularity value.

### Modular expression pattern of the AMPK and HIF-1 downstream genes

There exists a strong anti-correlation between the AMPK activity and HIF-1 activity across all 371 samples (Fig. 1A). In addition, expression of individual AMPK downstream genes was highly positively correlated within the AMPK gene group and negatively correlated with the HIF-1 downstream genes, and *vice versa* (Fig. 1B). The expression pattern of these genes was highly modular and consisted of two modules, one containing mainly AMPK-downstream genes and the other HIF-1-downstream genes, as identified by the Newman algorithm (Fig. 1C).

### Modularity and metabolism phenotypes

To evaluate the modular gene expression pattern of different metabolism phenotypes of HCC samples, we performed principal component analysis (PCA) on the RNA-Seq data of 33 AMPK downstream genes and 23 HIF downstream genes. Since AMPK and HIF-1 are master regulators of OXPHOS and glycolysis, respectively [19], the resulting first principal components (PC1s) for AMPK and HIF-1 downstream genes were assigned as the axes to quantify the activities of OXPHOS and glycolysis. After projecting all 371 HCC samples to the AMPK and HIF-1 axes, each HCC sample was assigned a metabolic state of glycolysis (HIF-1*^high^*/AMPK*^low^*), hybrid (HIF-1*^high^*/AMPK*^high^*) or OXPHOS (HIF-1*^low^*/AMPK*^high^*) through *k*-means clustering using the sum of absolute differences (Fig. 2A). Group modularity calculation showed that the OXPHOS group had the lowest mean modularity and glycolysis group had the highest mean modularity (Fig. 2B). Combined with survival curves of the three groups (Fig. 2C), it is clear that the glycolysis group had the worst survival and OXPHOS the best, with hybrid in the middle, indicating that higher modularity corresponded to a more aggressive tumor. In Fig. 2B and for all bar plots below, the error bars are obtained through the bootstrapping method. To obtain the significance levels, we used the method described in the ‘Materials and Methods’ section.

**Figure 2.**
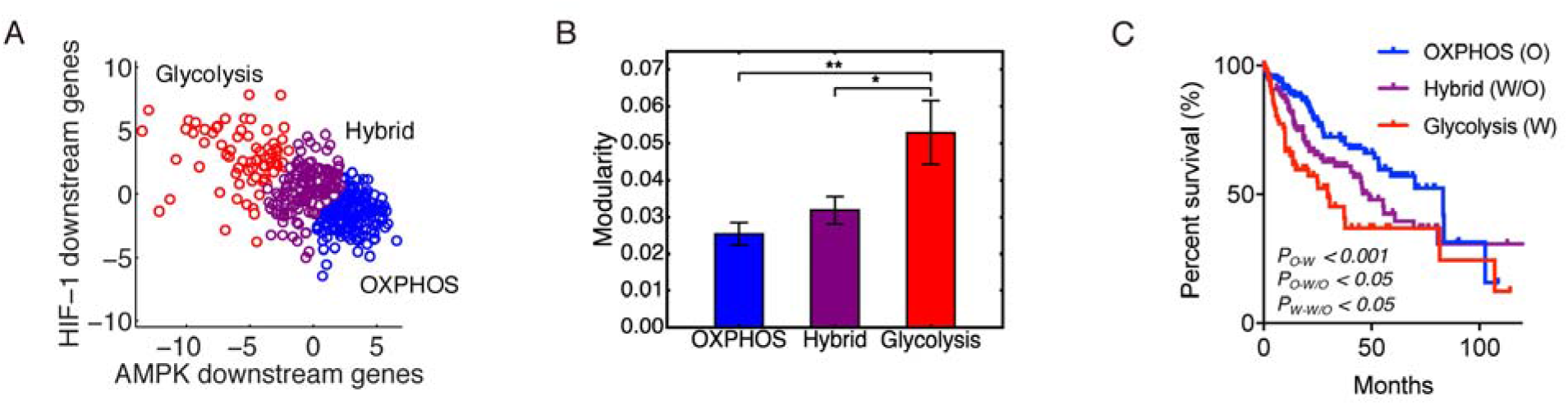
Modularity and metabolism phenotypes. **(A)** The 371 patients’ samples are clustered into three metabolism phenotypes: OXPHOS (blue), hybrid (magenta), and glycolysis (red). **(B)** Group modularity of three metabolism phenotypes. Here, ‘*’ represents 0.01 < *p* ≤ 0.05, and ‘**’ represents. If there is no labeling of the significance level, it means the difference is not significant. **(C)** Kaplan-Meier overall survival curves of HCC patients in OXPHOS, hybrid and glycolysis.

### HCC samples at later tumor stage have higher modularity

To analyze the change of modularity with respect to tumor stage, we classified the 348 of the 371 HCC samples that have neoplasm disease stage information into two groups, stage I (171 samples) and stage II-IV (177 samples). This was done to ensure that each group has similar number of samples. Group modularity calculations show that the HCC samples in the stage II-IV group had a significantly higher mean modularity than the HCC samples in the stage I group (Fig. 3A). HCC samples at stage II-IV had a significantly worse survival than samples at stage I (Fig. 3B), which further confirmed that higher modularity corresponded to worse survival, i.e. a more aggressive tumor.

**Figure 3.**
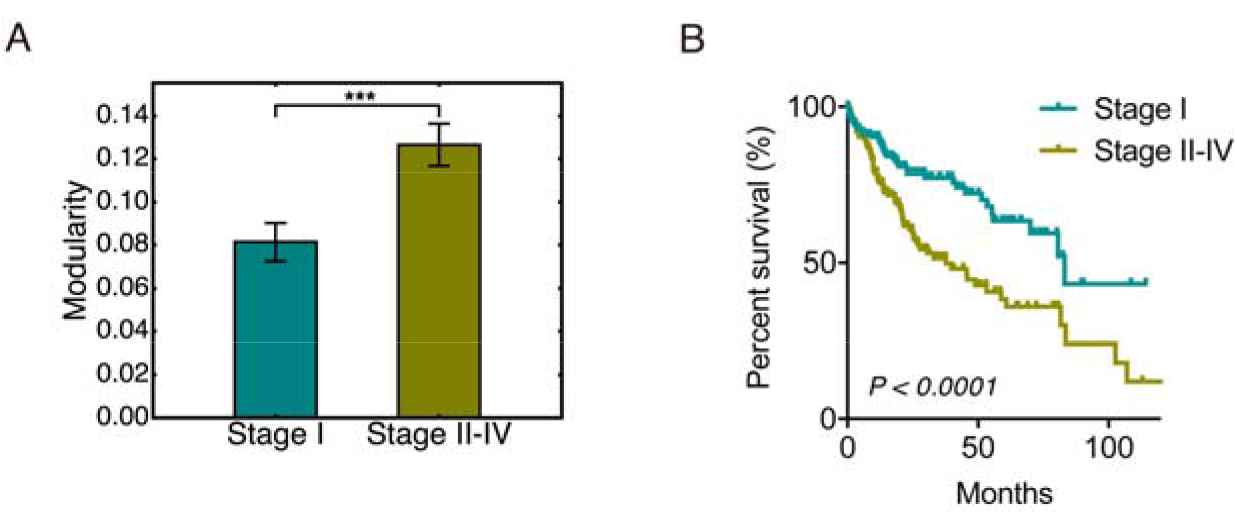
Modularity and tumor stages. **(A)** Bar plot of the group modularity of HCC patients at stage I and that at stage II-IV. Here ‘***’ represents *p* ≤ 0.001. **(B)** Kaplan-Meier overall survival curves of HCC patients at stage I and stage II-IV.

### HCC samples with higher metastatic potential have greater modularity

Metastasis accounts for more than 90% of cancer related deaths [26]. To evaluate the correspondence of modularity to metastatic potential of HCC samples, we grouped the samples based on their metastatic potential and calculated the group modularity. Genes SNRPF, EIF4EL3, HNRPAB, DHPS, PTTG1, COL1A1, COL1A2, LMNB1 (comprising the eight-gene signature) have been shown to be upregulated in metastases compared to primary tumor sites [27]. Expression levels of these genes has been used to evaluate the metastatic potential of primary tumors [27]. We here used the sum of *log*2-transformed values of expression levels of these eight genes to represent the metastatic potential of primary HCC samples. The 123 samples with the lowest metastatic potential were classified as the low potential group, and the 123 samples with the highest metastatic potential as the high potential group. Group modularity calculation results show that the high metastatic potential group had higher modularity and worse prognosis (Fig. 4A). We also used the expression of gene SPP1 to quantify the metastatic potential of HCC samples since the single SPP1 gene has been shown to be a diagnostic marker for metastatic HCC [28]. The grouping of HCC samples by expression of SPP1 show consistent results to that observed from the eight-gene signature (Fig. 4B). This result indicated that a highly modular pattern of cancer-associated gene interactions may serve as a sign of metastasis.

**Figure 4.**
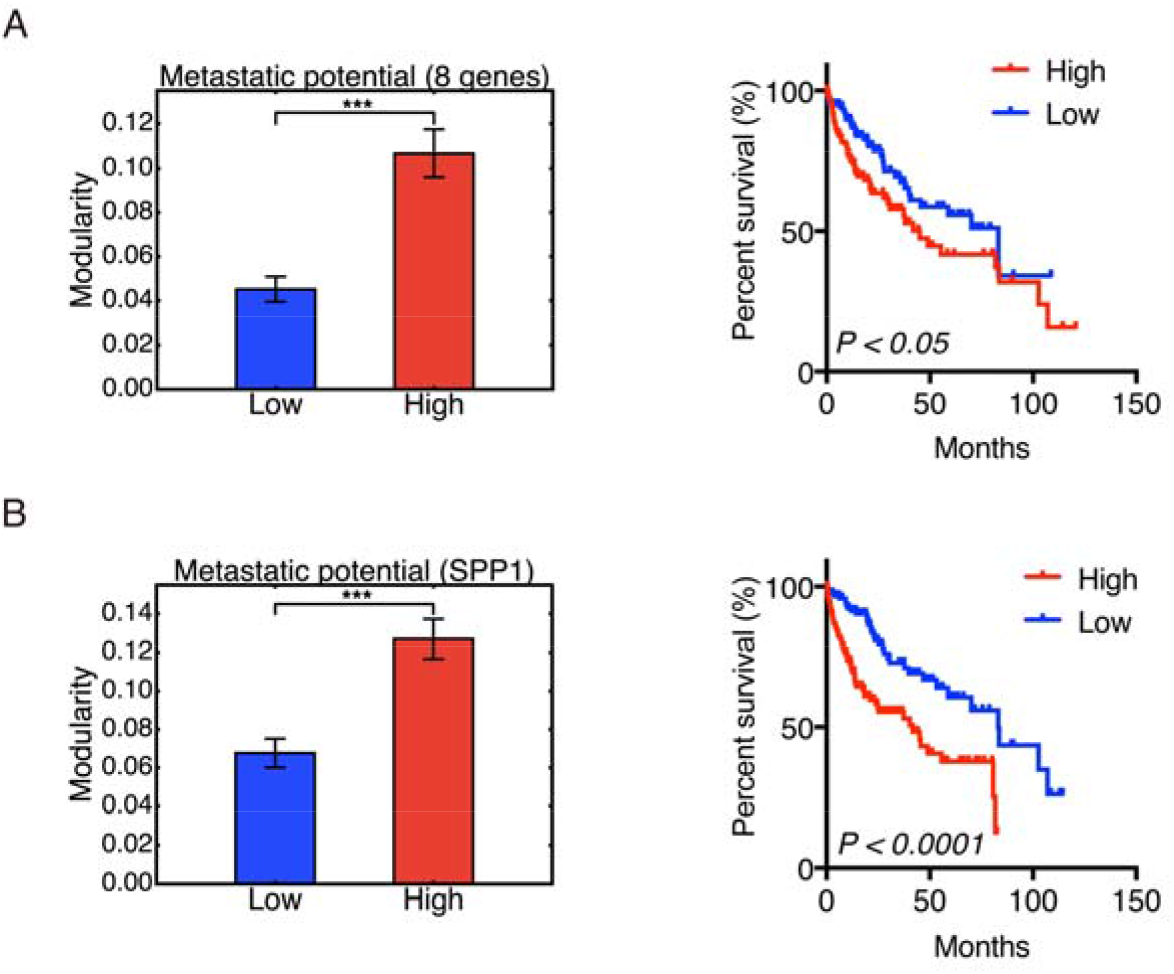
Modularity and metastatic potential. Left panel: Group modularity of HCC samples with low and high metastatic potential evaluated by eight-gene signature and SPP1 **(B)**. Right panel: Kaplan-Meier overall survival curves of HCC patients with low and high metastatic potential evaluated by eight-gene signature **(A)** and SPP1 expression **(B)**. Here ‘***’ represents *p* ≤ 0.001.

### Modularity and tumor recurrence

Tumor relapse is a supreme clinical challenge [29]. To analyze how tumor relapse is connected to the modularity of metabolic genes in HCC samples, we classified the 319 of 371 HCC samples that have tumor recurrence information – ‘recurred’ or ‘disease free’. Here the 319 samples were classified into non-recurrence and recurrence groups within 12 months, 24 months, or 36 months respectively. For example, the recurrence group within 12 months includes HCC samples whose disease-free status was ‘recurred’ and the ‘disease free time’ was shorter than 12 months. The non-recurrence group within 12 months includes HCC samples whose ‘disease free time’ was longer than 12 months, with either ‘recurred’ or ‘disease free’ status.

In all three cases, we observed that the group of HCC patients with recurred tumors had a higher mean modularity than the group of patients without recurred tumors (Fig. 5A). The difference between the recurrence and no-recurrence groups became more significant as time increased from 12 to 24 to 36 months. The survival curves confirmed that the recurrence group, which was also the high modularity group, had poor survival (Fig. 5A).

**Figure 5.**
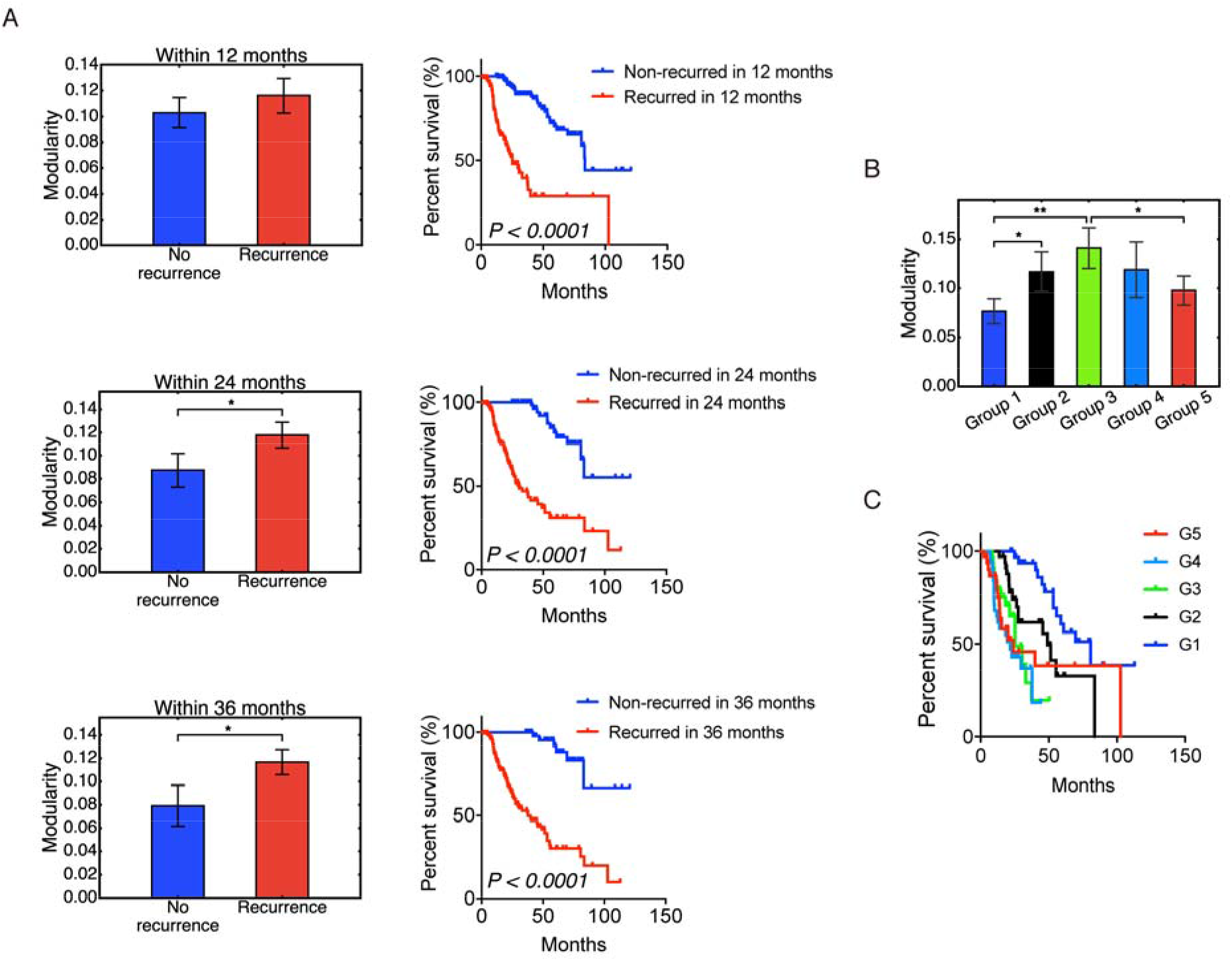
Modularity and tumor recurrence. **(A)** Modularity (left panels) and Kaplan-Meier overall survival curves (right panels) of patients that were stratified into recurrence and non-recurrence within 12, 24 and 36 months. **(B)** Non-monotonic change of modularity with tumor recurrence time. Samples in group 1 have the longest recurrence time and samples in group 5 has the shortest recurrence time. **(C)** Kaplan-Meier overall survival curves of group 1-5. Here ‘*’ represents 0.01 < *p* ≤ 0.05, and ‘**’ represents. If there is no labeling of the significance level, it means the difference is not significant. Significant p-values in **(C)** are as follows: *p*(*G*_1_, *G*_2_) < *p*(*G*_1_, *G*_4_) < 0.0001, *p*(*G*_1_, *G*_5_) < 0.01, *p*(*G*_2_, *G*_3_) < 0.05, *p*(*G*_2_, *G*_4_) < 0.01.

To understand the origin of the correlation between higher modularity and worse survival, we examined the relation between modularity and tumor recurrence time among recurred patients. Among the 319 samples, 174 have disease-free status as ‘recurred’. After discarding the 4 patients with the longest disease free time, the rest were sorted based on the disease-free time and classified into 5 groups – group 1, 2, 3, 4 and 5 with decreasing disease-free time. That is, group 1 had the longest recurrence time, and group 5 had the shortest recurrence time. The result is shown in Fig. 5B, and the corresponding survival curves for each group are shown in Fig. 5C. Modularity first increased with tumor progression, and then decreased. Even though the differences between each group were not always significant, the significant difference between group 1 and group 5 strongly supported this non-monotonic trend. It is also worth noting that modularity correlated with worse survival for the first 3 groups, but the correlation is reversed for groups 4 and 5. This result is similar to the trend observed in a study of acute myeloid leukemia [9]. At early stages, increased modularity correlates with decreased survival as cancer cells organize their gene expression against the host. At later stages, cancer has overcome the host defenses, and a high value of modularity is no longer selected for. Host survival, while low, becomes independent of modularity. We note that this crossover occurs rather late: recurrence times for groups 1, 2, 3 were 90-22 months, 22-13 months, and 13-8 months; the recurrence times for groups 4 and 5 were 8-4 and 4-1 months, respectively. Note that Fig. 5B and 5C are based on patients with recurred tumors only, whereas Fig. 5A contains both recurred patients and disease-free patients.

### Clinical application of modularity: Individual modularity and prediction

Calculation of group modularity is useful for understanding the group differences of metabolic gene expression patterns and the general relation between modularity and malignancy. However, for clinical application, individual modularity is required in order to make predictions regarding individual prognosis. The detailed definition and calculation procedure of individual modularity can be found in the ‘Materials and Methods’ section. Simply put, we applied the Newman algorithm to an individual cancer-associated gene network, with a new method to define links and with an additional de-noising step.

Individual modularity for all 371 samples ranged from 0.248 to 0.652, with mean 0.453 and standard deviation 0.079. These numbers appeared to be consistent with modularity values found in other functional human biological networks [30]. Modularity at the individual level largely confirmed the above group-level trends of modularity for HCC patients classified by metabolism phenotypes, stage information, recurrence status and metastatic potential. Higher individual modularity corresponded to the glycolysis phenotype, Fig. 6A, later tumor stage, Fig. 6B, tumor recurrence, Fig. S2A-C, and higher metastatic potential, as determined by the eight-gene signature, Fig. 6C, and SPP1 expression, Fig. S2D and Fig. S3. Together, these results validate the use of the metric of individual modularity to evaluate the aggressiveness of individual HCC patients.

**Figure 6.**
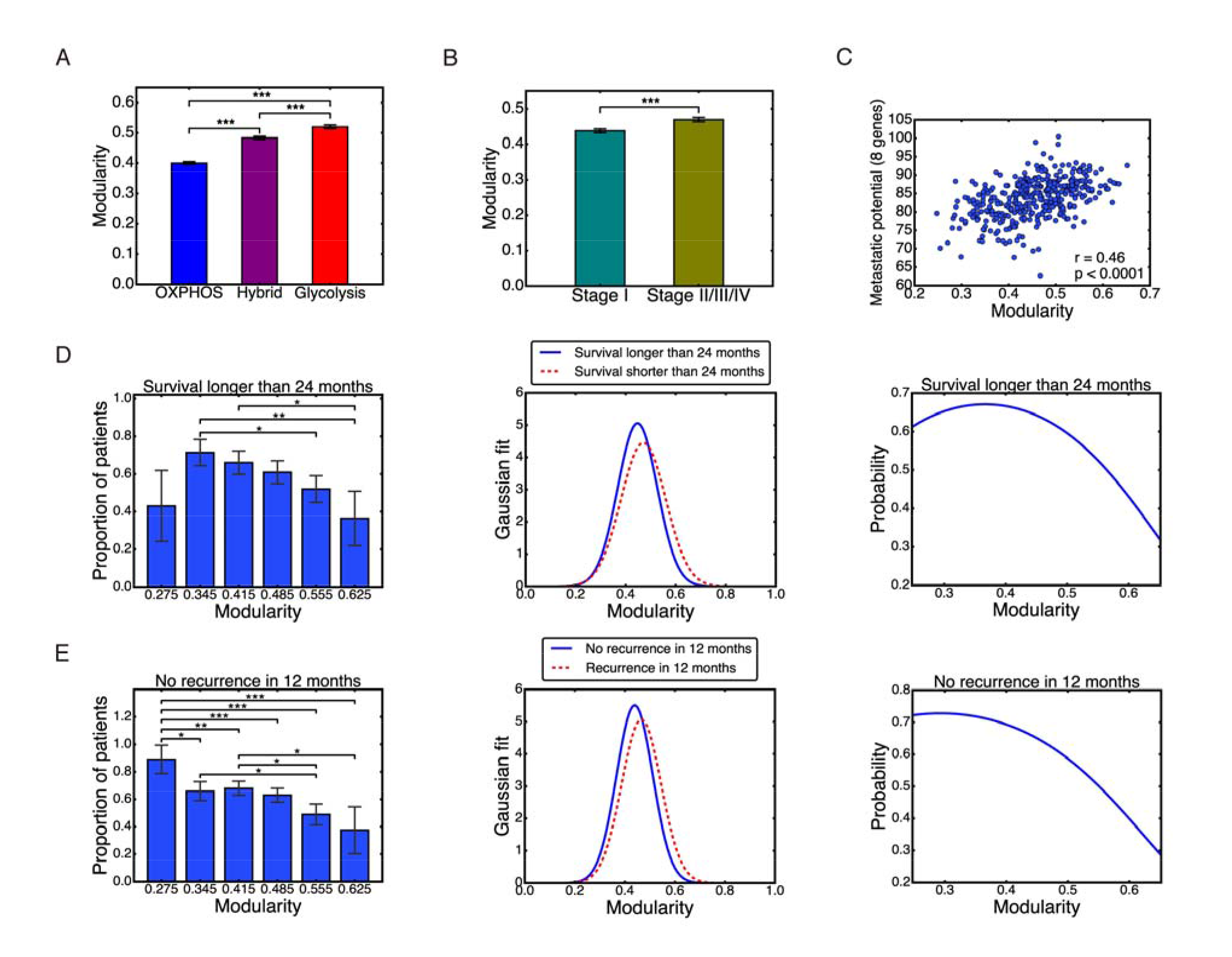
Individual modularity. **(A-C)** Individual modularity results show the same trend of modularity with metabolism types, stages, and metastatic potential. Pearson correlation between individual modularity and eight-gene metastatic potential r=0.46, p<0.0001. **(D)** Left: probability of survival longer than 24 months derived from data. Middle: Gaussian distribution of modularity values for the two groups. Right: same probability based on Gaussian model. **(E)** Left: probability of no recurrence in 12 months derived from data. Middle: Gaussian distribution of modularity values for the two groups. Right: same probability based on Gaussian model. Here ‘*’ represents 0.01 < *p* < 0.05, ‘**’ represents 0.001 < *p* <, and ‘***’ represents *p* ≤ 0.001. If there is no labeling of the significance level, it means the difference is not significant.

To make prognostic predictions with individual modularity, we focus on patients’ survival and tumor recurrence. We attempted to predict the probability of survival longer than 24 months and no recurrence in 12 months, so that each group has a comparable amount of samples: survived longer than 24 months, 140 samples; shorter than 24 months, 91 samples; no recurrence in 12 months, 176 samples; and recurrence within 12 months, 104 samples. We then divided patients into 6 groups based on their individual modularity values: 0.24-0.31, 0.31-0.38, 0.38-0.45, 0.45-0.52, 0.52-0.59, and 0.59-0.66. For each group, we counted the number of patients that survived longer 24 months and that remained disease-free for more than 12 months. We then calculated the proportion of these patients in each group, Fig. 6D-E, left panel. Overall, the higher the modularity, the lower the survival and disease-free probability. The only exception is the first bar in Fig. 6D left panel, which could potentially due to the very small number of 7 patients in the group.

We then captured these results by a Gaussian model of the modularity distribution of each group, with mean and standard deviation computed from individual modularity of each group, Fig. 6D and 6E, middle panel. Based on (eq.4) and (eq. 5) defined in the ‘Materials and Methods’ section, the probability of survival over 24 months and the probability of no recurrence in 12 months was calculated, Fig. 6D and 6E, right panel. This simple model was able to recapitulate the trend observed in the clinical data, Fig. 6D and 6E, left panel. The modularity range in these two plots was selected as 0.248 – 0.652 to match with the observed individual modularity values. Individual modularity showed significant potential as a predictor of patients’ survival or tumor recurrence. A high value of individual modularity was predictive of poor prognosis, with values of *M* > 0.6 correlated to survival and non-recurrence probabilities less than 0.4.

## Discussion

Metabolic reprogramming is an emerging hallmark of cancer [10, 11]. Both aerobic glycolysis and oxidative phosphorylation (OXPHOS) play important roles in orchestrating cancer metabolism [12-15, 17, 18, 31]. Previously, Yu *et al.* developed the AMPK and HIF-1 signatures to quantify the activities of metabolism phenotypes in hepatocellular carcinoma (HCC) [19]. There was a visually apparent modular pattern of gene expression due to the strong anti-correlation between AMPK and HIF-1 activities in HCC. In this work, we analyzed the gene expression pattern of metabolic genes in HCC in term of modularity and studied its correlation with metabolism phenotypes, tumor stages, metastatic potentials and tumor recurrence.

The analyses of modularity in the glycolysis, hybrid and OXPHOS metabolism phenotypes; stage I and stage II-IV tumor stages; and varying tumor metastatic potentials and recurrence status consistently showed that a higher modularity of the AMPK and HIF-1 downstream gene network corresponded to worse overall survival results of HCC patients. For example, a group of samples characterized by high glycolytic activity showed significantly higher modularity than a group of samples characterized by high OXPHOS activity, and worse prognosis. The result is consistent with the experimental observation that hepatocarcinogenesis initiates with a switch of metabolism from OXPHOS to glycolysis, and glycolysis is maintained to facilitate the aggressive features of advanced HCCs [32, 33]. Similarly, comparison of HCC samples at stage I to that at stage II-IV showed that HCC samples at stage II-IV have a more modular expression pattern of metabolic genes and worse survival prognosis. Additionally, HCC patients with a higher metastatic potential had a more modular expression pattern of metabolic genes and worse survival prognosis. Finally, patients with tumor recurrence within a given time had a higher modularity of the metabolic gene network and worse prognosis than patients with no tumor recurrence.

One interesting phenomena in this work is the non-monotonic relation between modularity and tumor progression as shown in Fig. 5B. Modularity increased first and then decreased. This result is similar to the trend observed in a previous study of acute myeloid leukemia [9]. We argue that at early stages of tumor progression, a modular pattern of cancer-associated gene interactions is organized by tumor cells, so that they can counteract the host defense systems. At later stages of tumor progression, cancer has overcome the host defenses, and a high value of modularity is no longer selected for. The results here suggest that the relation between modularity and tumor aggressiveness is mediated by tumor progression. For most of the patient’s history, a higher modularity indicates higher risk. Only when tumor progression has reached a very late stage, may a lower modularity indicate higher risk. Therefore, an accurate interpretation of modularity should take progression stage into consideration.

We further investigated the relation between modularity and tumor recurrence time in three subsets: HCC samples with glycolysis phenotype, at stage II-IV, and with high metastatic potential determined by the eight-gene signature (Fig. S4). These groups were chosen as they tend to be under highly stressful conditions such as hypoxia due to rapid proliferation of tumor and response from the host immune systems during metastasis. The glycolysis group had 37 patients that recurred. After discarding 2 samples with the shortest recurrence time, the rest were distributed into 5 equal size groups. A similar procedure was taken for the other two groups. Again, HCC samples in group 1 had the longest recurrence time and HCC samples in group 5 had the shortest recurrence time. Survival curves for each group were also plotted. We found that the correlation of higher modularity with worse prognosis exists for the roughly ~60% (top 3 groups) of patients with the longest recurrence time in all three cases. Interestingly, a reversal of this correlation occurs at about the same recurrence time: 9.1 months for the glycolysis group, 6.4 months for the stage II-IV group, and 7.9 months for the high metastatic potential group. These times are consistent with the reversal of the correlation at 8 months found among all recurred patients (Fig. 5B).

Taken together, these results show that modularity is selected for under the stressful conditions of early to mid-progression. That is, more aggressive early and mid-progression tumors, as judged by decreased host survival probability, have higher modularity of metabolic genes. These results confirm our previous hypothesis that more malignant tumors are usually characterized by a more modular expression pattern of cancer-associated genes [8, 9]. We predict that higher modularity increases the fitness of tumors because metabolic networks are typically under increased stress in HCC tumor cells [34]. Thus, tumors with a more modular metabolic gene network typically are more fit and are more likely to overcome the body’s defenses. Once the transition to imminent recurrence is achieved, the selection strength for modularity is no longer present, and the observed values of modularity decrease.

Notably, modularity is predictive of prognosis independent of metastatic status of HCC samples. We analyzed the association of modularity with different metabolism phenotypes, varying stages, and tumor recurrence for the HCC patients with no distant metastasis, cancer staging ‘M0’, i.e. no spread of tumor to other parts of the body, Fig. S5. More modular gene expression patterns of metabolic genes were observed for HCC samples in the glycolysis phenotype than in the OXPHOS phenotype, Fig. S6, at stage II-IV than that at stage I, Fig. S7, and with tumor recurrence than without recurrence, Fig. S8. These results support the conclusion that modularity is a fundamental order parameter correlated with tumor aggressiveness.

To the best of our knowledge, this is the first effort to evaluate aggressiveness of HCC samples by evaluating the expression pattern of metabolic genes in terms of modularity. Further work can extend the modularity concept to different types of tumors. There are at least two avenues for the improvement of the present study. First, a different set of parameters used in calculating individual modularity might affect the predictive efficiency. We list in Table S2 the parameters for calculating individual modularity using ITSPCA. Varying values of these standard parameter gave similar results, but with a weaker signal. We therefore believe that the chosen parameter set works well and keeps most of the signal. Future work could quantify the signal as a function of the parameter set to improve the predictive power of individual modularity. Second, temporal expression profiles of the metabolic genes in HCC samples from individual patients may further power the personalized prognosis.

In summary, modular interactions between metabolic genes in HCC play a key role in HCC prognosis. HCC patients with higher Individual modularity have a higher risk of tumor recurrence and poorer prognosis. There are several possible clinical applications from the individual modularity. First, prediction of patient survival and recurrence probabilities with individual modularity can adjust the choice of appropriate therapies. Second, key driver genes promoting HCC progression could be potentially identified, *e.g.* a hub node gene that strengthens intracommunity interactions and increases modularity. Third, drug treatment efficacy could be evaluated by testing the ability of drugs to disrupt the modular interactions between cancer-associated genes. A novel approach for drug design could target genes that significantly contribute to the increase of modularity of the cancer-associated gene network.

## Materials and Methods

### 1. 371 primary HCC samples

RNA-Seq data for 373 hepatocellular carcinoma (HCC) samples, which contain the gene expression of 33 AMPK downstream genes and 23 HIF-1 downstream genes, were obtained from TCGA at cBioPortal [35, 36]. Among the 373 HCC samples, 371 primary tumor samples were used for subsequent analysis, and 2 recurrent tumor samples were excluded.

### 2. Calculation of Group Modularity

Modularity of a given graph *A_ij_* was defined as

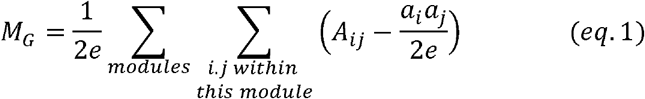

where *A_ij_* is 1 if there is an edge between nodes *i* and *j* and 0 otherwise, the value of *a_i_* = Σ*_j_ A_j_* is the degree of node *i*, and e = ½ Σ*_i_a_i_* is the total number of edges. This definition can be extended to unsigned weighted graphs, where *A_ij_* is the weight of the edge between nodes *i* and *j* and where *A_ij_* > 0. Here the subscript ‘G’ is used because this definition is adopted for calculation of group modularity. We applied Newman’s algorithm [21] to graph *A_ij_* to calculate modularity. This algorithm found the partition of 56 genes into modules that maximized modularity *M_G_*. This maximized modularity was used as the final modularity value for data analysis.

To calculate modularity of HCC samples grouped by metabolism phenotypes, tumor stages, metastatic potential, or recurrence status, the RNA-seq data of each of the 56 AMPK and HIF-1 downstream genes the gene expression data were transformed by log_2_ and normalized, i.e.

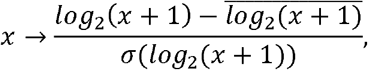

where *x* represents the expression of each gene, 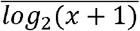 is the mean of the *log*_2_ transformed values across all patients’ expression of this gene, and *σ*(*log*_2_(*x* + 1)) is the standard deviation of the *log_2_* transformed values.

The metabolic gene network for each group was defined by setting the 56 genes as the nodes and the Pearson correlation coefficient between genes as the link weights. The resulting network was represented by a 56*56 correlation matrix *C.* Since the above definition of modularity is for an unsigned graph, and since we regard negative correlations as weak links between genes, the whole matrix was shifted as *C′* = (*C* + 1). We then set the diagonal elements of to 0 to eliminate self-loops and used the Newman algorithm to calculate the modularity of this matrix *C′*.

To compare modularity between different patient groups, *e.g.* glycolysis versus OXPHOS, the bootstrapping method was used. This method takes the observed individual gene expression values as the most representative measure of the underlying distribution of expression values. That is, the distribution of expression values is taken as a sum over *δ* functions at the observed values. Predictions are computed from samples taken from this estimated distribution. For example, for the glycolysis group of 75 patients, the gene expression correlation matrix was calculated by randomly taking expression values from the 75 patients with replacement. The modularity of this correlation matrix was computed as described above. This sampling process was repeated 10 000 times to obtain 10 000 modularity values for the glycolysis group. Mean modularity and standard error were then obtained. This same procedure was used to compute modularity for each of the other groups. Calculation of p-values is described in subsection 6.

### 3. Calculation of Individual Modularity

Typically, for each patient there is one expression value for each gene, and no correlation between genes based upon only a single patient’s data can be computed. We propose, therefore, to define the link between gene *i* and gene *j* of patient *α* as

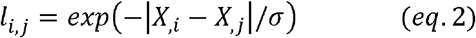

where *X_,i_* is the expression of gene *i* of patient □ and is the standard deviation of |*X_,i_* – *X_,i_*| averaged across all pairs of genes and all patients, with *σ* = 57 887 in our case. This definition considers the link between gene *i* and gene *j* weak if the distance between them, i.e.|*X_,i_* – *X_,i_*|, is large. The scaling by *σ* ensures that |*X_,i_* – *X_,i_*|/*σ* remains within a reasonable order of magnitude.

Unlike the group modularity case, having only 56 expression values for each individual means the noise in the data has a greater impact on the calculated modularity values. Thus, a better way of filtering noise is needed. A standard approach is to reconstruct the data based only on cleaned leading eigenvectors. We utilized the iterative thresholding sparse PCA (ITSPCA) algorithm for this purpose [37]. The algorithm starts by keeping only the top eigenvectors. To separate signal and noise, such that signal is defined as above a threshold, a wavelet transformation is used, see Supplementary Materials section 2, Fig. S1 and Table S1. Data that were dense in real space became sparse in wavelet space, and a cutoff was then applied in wavelet space to eliminate the noise. The standard wavelet transformation algorithm requires that the number of entries be a power of two. Zero-padding was applied to the input data matrix *X* so that it became a 371*64 matrix, with the last 8 columns containing only zeros. The ITSPCA algorithm output the cleaned 56* *n* matrix version of the leading eigenvectors *P_n_*, which is used to reconstruct the raw data as

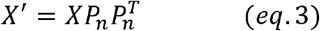

where *X* is the original raw data matrix, and *X’* is the reconstructed matrix that has the same dimension of *X*. The cleaned data *X’* should contain mostly signal and much less noise than *X*, and therefore *X’* was used in calculation of links (eq.2). Note that, unlike the group modularity calculation, is based on the raw data without taking a logarithm. This is because we believe that noise had already been filtered out by ITSPCA, and taking the logarithm would only weaken the signal. See Table S2 for chosen input parameters of the ITSPCA algorithm.

After determining *X’*, we computed the individual gene network linkage based on (eq.2). We then applied the binarization step where the top 5.6% edges (178 edges) were set to 1 and the rest set to zero. According to our previous work [30], this binarization step increases the signal-to-noise ratio without discarding important information. The Newman algorithm was used to compute modularity for each patient, *M_i_*. We bootstrapped the individual modularity 10,000 times for each group, and we calculated the mean of the 10,000 individual modularity. The average and standard deviation of the means were plotted in the bar plots. Note that this average is the same as the one directly calculated from the vector of *M_i_* and this standard deviation is the same as the standard error of the mean directly calculated from the vector of *M_i_*. P-values were calculated using the method described in subsection 6.

### 4. Definition of probability of surviving longer than 24 months based on individual modularity

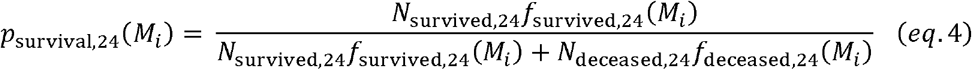

where *N*_survived,24_ and *N*_decreased,24_ are the numbers of patients that lived longer than 24 months and deceased within 24 months, respectively. Here *f*_survived,24_ and *f*_decreased,24_ are the probability density functions of the modularity distribution of survived and deceased group, respectively. Given modularity *M_i_* we calculated *p*_survival,24_ and thus obtained the probability curve of surviving more than 24 months.

### 5. Definition of probability of no recurrence in 12 months based on individual modularity

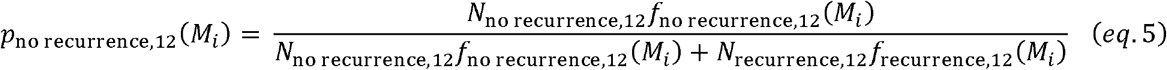

where *N*_no recurrence,12_ and *N*_recurrence,12_ are the numbers of patients that remained disease free for more than 12 months and those that recurred within 12 months, respectively. Here *f*_no recurrence,12_ and *f*_recurrence,12_ are the probability density functions of the modularity distribution of disease-free and recurred group, respectively. Given modularity *M_i_*, we calculated *p*_no recurrence,12_ and thus obtained the probability curve of no recurrence within 12 months.

### 6. *P*-value calculation

Given two samples *x*_1_ and *x*_2_, a standard way to test for equal means is a two-sample *t* test. However, in the current research, it is often the case that we do not have direct access to *x*_1_ and *x*_2_, or that the original *x_1_* and *x_2_* are of no interest. For example, in the case of comparing group modularity, *e.g.* Fig. 2B, 3A, the only available values are vectors of the bootstrapped modularity of each group. The original *x_1_* and *x*_2_, which are the gene expression of samples in the group, were of no interest.

We therefore perform a standard Monte Carlo test of *p*-values. Given input data *x*_1_, *x*_2_, and function of interest *F*, we perform *B* bootstrap samples of the function with replacement, obtaining vectors F_1_ and F_2_. Each element of F_i_ was obtained by calculating 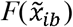, where 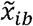 is the bootstrapped sample of *x_i_* in bootstrap *b* (*b*=1,2, …, B). We assume the average of F_1_ is greater than F_2_. We define

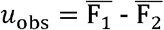

and shift the bootstrap samples as

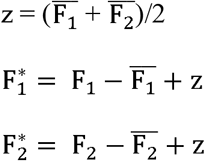

We define for each bootstrap sample *b*

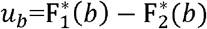

Then the *p*-value is defined as

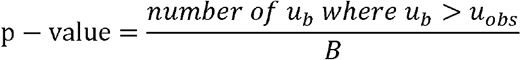

The bootstrapping process gives the distribution of F_1_ and F_2_. Note that z is the mean of the concatenated vector of F_1_ and F_2_, and B is set to 10 000 in all cases. The null hypothesis H_0_ is that F_1_ and F_2_ have equal means, and the alternative hypothesis H_1_ is that the mean of F_1_ is larger than that of F_2_. We have confirmed that, if function 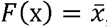, Monte Carlo test gives the same *p*-values as one-tailed *t* test on (*x*_1_, *x*_2_).

For bar plots involving group modularity, *x*_1_ and *x*_2_ are gene expressions of group 1 and group 2, and *F* is the modularity. For bar plots involving individual modularity, *x*_1_ and *x*_2_ are individual modularity of group 1 and group 2, and *F* is the mean. For bar plots involving proportion of patients, *x*_1_ and *x*_2_ are disease free time or survival time of group 1 and group 2, and *F* is the proportion of patients that were disease-free for more than 12 months, of survived longer than 24 months.

## References

1. Byam J, Renz J, M¡ll¡s JM. Liver transplantation for hepatocellular carcinoma. Hepatobiliary Surg Nutr. 2013; 2: 22–30. doi: 10.3978/j.issn.2304-3881.2012.11.03.

2. Jemal A, Bray F, Center MM, Ferlay J, Ward E, Forman D. Global cancer statistics. CA Cancer J Clin. 2011; 61: 69–90. doi: 10.3322/caac.20107.

3. Raza A, Sood GK. Hepatocellular carcinoma review: current treatment, and evidence-based medicine. World J Gastroenterol. 2014; 20: 4115–27. doi: 10.3748/wjg.v20.il5.4115.

4. Lou J, Zhang L, Lv S, Zhang C, Jiang S. Biomarkers for Hepatocellular Carcinoma. Biomark Cancer. 2017; 9:1-9. doi: 10.1177/1179299X16684640.

5. Davila JA, Morgan RO, Shaib Y, McGlynn KA, El-Serag HB. Diabetes increases the risk of hepatocellular carcinoma in the United States: a population based case control study. Gut. 2005; 54: 533–9. doi: 10.1136/gut.2004.052167.

6. Creixell P, Reimand J, Haider S, Wu G, Shibata T, Vazquez M, Mustonen V, Gonzalez-Perez A, Pearson J, Sander C, Raphael BJ, Marks DS, Ouellette BFF, et al. Pathway and network analysis of cancer genomes. Nat Methods. 2015; 12: 615–21. doi: 10.1038/nmeth.3440.

7. Serra-Musach J, Mateo F, Capdevila-Busquets E, de Garibay GR, Zhang X, Guha R, Thomas CJ, Grueso J, Villanueva A, Jaeger S, Heyn H, Vizoso M, Perez H, et al. Cancer network activity associated with therapeutic response and synergism. Genome Med. 2016; 8: 88. doi: 10.1186/s13073-016-0340-x.

8. Chen M, Deem MW. Hierarchy of gene expression data is predictive of future breast cancer outcome. Physical Biology. 2013; 10: 056006. doi: 10.1088/1478-3975/10/5/056006.

9. Tripathi S, Deem MW. Hierarchy in gene expression is predictive of risk, progression, and outcome in adult acute myeloid leukemia. Physical Biology. 2015; 12: 016016. doi: 10.1088/1478-3975/12/1/016016.

10. Hanahan D, Weinberg RA. Hallmarks of cancer: the next generation. Cell. 2011; 144: 646–74. doi: 10.1016/j.cell.2011.02.013.

11. Pavlova NN, Thompson CB. The Emerging Hallmarks of Cancer Metabolism. Cell Metab. 2016; 23: 27–47. doi: 10.1016/j.cmet.2015.12.006.

12. Warburg O. On the origin of cancer cells. Science. 1956; 123: 309–14. doi: 10.1126/science. 123.3191.309.

13. Vander Heiden MG, Cantley LC, Thompson CB. Understanding the Warburg effect: the metabolic requirements of cell proliferation. Science. 2009; 324:1029–33. doi: 10.1126/science. 1160809.

14. Lu CL, Qin L, Liu HC, Candas D, Fan M, Li JJ. Tumor cells switch to mitochondrial oxidative phosphorylation under radiation via mTOR-mediated hexokinase II inhibition—a Warburg-reversing effect. PLoS One. 2015; 10: e0121046. doi: 10.1371/journal.pone.0121046.

15. Viale A, Pettazzoni P, Lyssiotis CA, Ying H, Sanchez N, Marchesini M, Carugo A, Green T, Seth S, Giuliani V, Kost-Alimova M, Muller F, Colla S, et al. Oncogene ablation-resistant pancreatic cancer cells depend on mitochondrial function. Nature. 2014; 514: 628–32. doi: 10.1038/naturel3611.

16. Tan AS, Baty JW, Dong LF, Bezawork-Geleta A, Endaya B, Goodwin J, Bajzikova M, Kovarova J, Peterka M, Yan B, Pesdar EA, Sobol M, Filimonenko A, et al. Mitochondrial genome acquisition restores respiratory function and tumorigenic potential of cancer cells without mitochondrial DNA. Cell Metab. 2015; 21: 81–94. doi: 10.1016/j.cmet.2014.12.003.

17. Maiuri MC, Kroemer G. Essential role for oxidative phosphorylation in cancer progression. Cell Metab. 2015; 21:11–2. doi: 10.1016/j.cmet.2014.12.013.

18. Porporato PE, Payen VL, Perez-Escuredo J, De Saedeleer CJ, Danhier P, Copetti T, Dhup S, Tardy M, Vazeille T, Bouzin C, Feron O, Michiels C, Gallez B, et al. A mitochondrial switch promotes tumor metastasis. Cell Rep. 2014; 8: 754–66. doi: 10.1016/j.celrep.2014.06.043.

19. Yu L, Lu M, Jia D, Ma J, Ben-Jacob E, Levine H, Kaipparettu BA, Onuchic JN. Modeling the Genetic Regulation of Cancer Metabolism: Interplay between Glycolysis and Oxidative Phosphorylation. Cancer Res. 2017; 77: 1564–74. doi: 10.1158/0008-5472.CAN-16-2074.

20. Girvan M, Newman ME. Community structure in social and biological networks. Proc Natl Acad Sci USA. 2002; 99: 7821–6. doi: 10.1073/pnas.122653799.

21. Newman ME. Modularity and community structure in networks. Proc Natl Acad Sci USA. 2006; 103: 8577–82. doi: 10.1073/pnas.0601602103.

22. Dunne JA, Williams RJ, Martinez ND. Food-web structure and network theory: The role of connectance and size. Proc Natl Acad Sci USA. 2002; 99:12917–22. doi: 10.1073/pnas.192407699.

23. Pandey J, Koyuturk M, Grama A. Functional characterization and topological modularity of molecular interaction networks. BMC Bioinformatics. 2010; 11 Suppl 1: S35. doi: 10.1186/1471-2105-11-S1-S35.

24. Sun J, Deem MW. Spontaneous emergence of modularity in a model of evolving individuals. Physical Review Letters. 2007; 99: 228107. doi: 10.1103/PhysRevLett.99.228107.

25. Deem MW. Statistical Mechanics of Modularity and Horizontal Gene Transfer. Annual Review of Condensed Matter Physics, Vol 4. 2013; 4: 287–311. doi: 10.1146/annurev-conmatphys-030212-184316.

26. Gupta GP, Massague J. Cancer metastasis: building a framework. Cell. 2006; 127: 679–95. doi: 10.1016/j.cell.2006.11.001.

27. Ramaswamy S, Ross KN, Lander ES, Golub TR. A molecular signature of metastasis in primary solid tumors. Nat Genet. 2003; 33: 49–54. doi: 10.1038/ng1060.

28. Ye QH, Qin LX, Forgues M, He P, Kim JW, Peng AC, Simon R, Li Y, Robles Al, Chen Y, Ma ZC, Wu ZQ, Ye SL, et al. Predicting hepatitis B virus-positive metastatic hepatocellular carcinomas using gene expression profiling and supervised machine learning. Nat Med. 2003; 9: 416–23. doi: 10.1038/nm843.

29. Mitra A, Mishra L, Li S. EMT, CTCs and CSCs in tumor relapse and drug-resistance. Oncotarget. 2015; 6:10697-711. doi: 10.18632/oncotarget.4037.

30. Yue Q, Martin R, Fischer-Baum S, Ramos-Nuñez Al, Ye F, Deem MW. Brain modularity mediates the relation of cognitive performance to task complexity. J Cogn Neurosci. 2017; 29: 1532–46. doi: 10.1162/jocn_a_01142.

31. Tan AS, Baty JW, Dong LF, Bezawork-Geleta A, Endaya B, Goodwin J, Bajzikova M, Kovarova J, Peterka M, Yan B, Pesdar EA, Sobol M, Filimonenko A, et al. Mitochondrial Genome Acquisition Restores Respiratory Function and Tumorigenic Potential of Cancer Cells without Mitochondrial DNA. Cell Metabolism. 2015; 21: 81–94. doi: 10.1016/j.cmet.2014.12.003.

32. Kowalik MA, Guzzo G, Morandi A, Perra A, Menegon S, Masgras I, Trevisan E, Angioni MM, Fornari F, Quagliata L, Ledda-Columbano GM, Gramantieri L, Terracciano L, et al. Metabolic reprogramming identifies the most aggressive lesions at early phases of hepatic carcinogenesis. Oncotarget. 2016; 7: 32375–93. doi: 10.18632/oncotarget.8632.

33. Yang J, Wang C, Zhao F, Luo X, Qin M, Arunachalam E, Ge Z, Wang N, Deng X, Jin G, Cong W, Qin W. Loss of FBP1 facilitates aggressive features of hepatocellular carcinoma cells through the Warburg effect. Carcinogenesis. 2017; 38:134-43. doi: 10.1093/carcin/bgw109.

34. Wang MD, Wu H, Huang S, Zhang HL, Qin CJ, Zhao LH, Fu GB, Zhou X, Wang XM, Tang L, Wen W, Yang W, Tang SH, et al. HBx regulates fatty acid oxidation to promote hepatocellular carcinoma survival during metabolic stress. Oncotarget. 2016; 7: 6711–26. doi: 10.18632/oncotarget.6817.

35. Cerami E, Gao J, Dogrusoz U, Gross BE, Sumer SO, Aksoy BA, Jacobsen A, Byrne CJ, Heuer ML, Larsson E, Antipin Y, Reva B, Goldberg AP, et al. The cB¡o cancer genomics portal: an open platform for exploring multidimensional cancer genomics data. Cancer Discov. 2012; 2: 401–4. doi: 10.1158/2159-8290.CD-12-0095.

36. Gao J, Aksoy BA, Dogrusoz U, Dresdner G, Gross B, Sumer SO, Sun Y, Jacobsen A, Sinha R, Larsson E, Cerami E, Sander C, Schultz N. Integrative analysis of complex cancer genomics and clinical profiles using the cB¡oPortal. Sci Signal. 2013; 6: p|1. doi: 10.1126/scisignal.2004088.

37. Ma ZM. Sparse Principal Component Analysis and Iterative Thresholding. Annals of Statistics. 2013; 41: 772–801. doi: 10.1214/13-Aosl097.

